# Yck2 links mitochondrial function and pH homeostasis to cell wall remodeling and antifungal susceptibility in *Cryptococcus neoformans*

**DOI:** 10.64898/2026.09.02.749035

**Authors:** Amanda LM Bloom, David Goich, Eric Mendelson, Shichen Chen, Maia Mazzaferro, Jun Qu, John C. Panepinto

**Author notes:** Amanda LM Bloom, Roles: Conceptualization, Data Curation, Formal Analysis, Investigation, Methodology, Project Administration, Supervision, Validation, Visualization, Writing – Original Draft Preparation. David Goich, Roles: Investigation, Data Curation, Formal Analysis. Eric Mendelson, Roles: Investigation, Formal Analysis, Validation. Shichen Chen:, Roles: Data Curation, Investigation. Maia Mazzaferro, Roles: Investigation, Formal Analysis, Validation. Jun Qu, Roles: Resources, Supervision. John C. Panepinto, Roles: Conceptualization, Funding Acquisition, Supervision.

## Abstract

Invasive fungal infections are on the rise due to climate change, antifungal resistance, and increased usage of immunomodulating therapies. The antifungal arsenal is limited in number and in efficacy necessitating novel therapeutics. Deletion of the gene encoding the fungal yeast casein kinase, *YCK2*, has pleiotropic effects, impacting morphology, drug resistance, metabolism, cell wall, and virulence. Yck2 has recently been shown to be druggable, however the mechanisms by which Yck2 elicits its impacts are unknown. We show that *C. neoformans* Yck2 is a regulator of cell wall masking, thermotolerance, general stress, and drug resistance. Proximity labeling of Yck2-interacting proteins revealed that Yck2 interacts with proteins that function in the mitochondria as well as the essential plasma membrane H+ ATPase, Pma1. A role for Yck2 in mitochondrial regulation was supported by sensitivity to mitochondrial inhibitors and increased mitochondrial ROS production in the absence of Yck2, and presence of Yck2 in mitochondrial fractions. Using pHluorin-expressing cells we also found that intracellular pH is elevated in cells lacking Yck2 supporting a role in regulating Pma1 activity. Both mitochondria and cellular pH may contribute to cellular signaling and may explain the multitude of phenotypes associated with the *yck2*Δ mutant. Our results begin to identify the mechanisms by which Yck2 contributes to fungal cellular homeostasis which may support efforts to optimize Yck2 inhibitors.

**Author Summary:** *Cryptococcus neoformans* is a pathogenic fungus that causes ∼20% of AIDS-related mortality. Current antifungals are associated with resistance, high-cost, and/or ineffectiveness necessitating the need for novel therapies. We found that the fungal casein kinase, Yck2, contributes to cell wall remodeling, thermotolerance, drug resistance, and stress response, all of which are important for fungal pathogenesis. Our work revealed that Yck2 modulates mitochondrial function and regulates the intracellular pH, both of which may have cascading effects on the cell. Recent work has shown that fungal Yck2 can be successfully targeted for inhibition. Our data begins to elucidate how Yck2 is functioning in the cell, supporting efforts to develop Yck2 inhibitors as antifungals.

## Introduction

Invasive fungal infections are increasing at an alarming rate and the need for novel drug targets for antifungal development is urgent. In 2022, the WHO named *Cryptococcus neoformans* the number one fungal priority pathogen, advocating for better surveillance, research and development, and public health intervention (1). *C. neoformans* is a basidiomycetous environmental fungus that causes cryptococcal meningitis in individuals with compromised immunity, accounting for 20% of AIDS related mortality and causing invasive disease in solid organ transplant recipients with mortality rates up to 50% (2, 3). Gold standard therapy consists of the combination of amphotericin B + 5-fluorocytosine, a costly regimen that is unavailable in most resource-limited areas and is still associated with 30% mortality (4, 5). The azole class of antifungals, namely fluconazole, are fungistatic and associated with resistance (6–8). *C. neoformans* is inherently resistant to the latest developed class of antifungals, the echinocandins, which target the cell wall β-glucan synthase, eliminating its use to treat cryptococcal disease.

The fungal cell wall contains several pathogen-associated molecular patterns (PAMPs), making regulators of cell wall modeling enticing targets for antifungal therapies. Changes in the cell wall in response to the environment provide protection, and in the context of a human host promotes evasion from immune cells (9, 10). Our previous work demonstrated that in response to host temperature stress the cell wall is remodeled, keeping the pathogen associated molecular pattern (PAMP), β-1,3-glucan, masked, and this remodeling is dependent on reprogramming the translatome (11). Kinases serve as hubs for cellular signaling and play important roles in regulation. Screening a kinase knock-out collection for β-1,3-glucan unmasking lead to our identification of the Hog1/p38 MAPK and Mpk1 MAPK signaling modules as key contributors of cell wall remodeling following host temperature stress (12).

In our screen for kinases that regulate cell wall masking we also identified the yeast casein kinase, Yck2. Yck2 is a type I casein kinase and homolog of the *Saccharomyces cerevisiae* homologs Yck1 and its paralog Yck2, which rose due to genome duplication. Yck1/2 homologs are palmitoylated plasma membrane kinases (13, 14). In *S. cerevisiae* Yck1/2 plays a role in glucose signaling, phosphorylating repressors of glucose-responsive genes to promote their degradation by the proteasome when glucose is present (15, 16). Yck1/2 also phosphorylates the essential plasma membrane ATPase Pma1 upon glucose depletion contributing to inhibition of its proton pumping activity (17). In the opportunistic fungal pathogen *Candida albicans*, Yck2 regulates hyphal growth, biofilm production, cell wall integrity and the ability to cause damage to host cells (18), however Yck2 targets have not been identified, and it is currently unknown how these cellular processes are specifically regulated via Yck2. Interestingly, treatment of *C. albicans* echinocandin-resistant strains with compounds shown to target and inhibit Yck2 restores echinocandin susceptibility (19). The Cowen group recently derivatized these inhibitors and demonstrated their enhanced in vivo stability, limited effect on human CK1, and strong specificity for *C. albicans* Yck2, successfully demonstrating that fungal Yck2 is druggable (20).

*C. neoformans* Yck2, previously referred to as Cck1, was found to impact Mpk1 and Hog1 signaling and promote cell wall integrity and virulence (21). Specific targets of *C. neoformans* Yck2 and its effect on gene expression have not been explored. We hypothesized that Yck2 governs cell wall remodeling in response to host temperature, aiding in host cell evasion, and set out to understand how Yck2 regulates this process. In addition to its important role in cell wall remodeling, we found that Yck2 is an important regulator of antifungal susceptibility and general stress. Our transcriptomics and interactome analyses strongly suggest that Yck2 regulates and interacts with mitochondria, validated by mitochondrial dysregulation in the *yck2*Δ mutant. Further, we identified the essential plasma membrane H+ ATPase, Pma1, as a 37°C dependent target of Yck2. Our work here suggests that Yck2 acts at both the plasma membrane and mitochondria to regulate drug sensitivity, cell wall remodeling and thermotolerance.

## Results

### Yck2 regulates cell wall masking and architecture at host temperature

In a screen to determine signaling pathways involved in cell wall remodeling in response to host temperature stress, we previously screened a *C. neoformans* kinase knock out collection for β-1,3-glucan unmasking by flow cytometry. From this screen we identified kinases in the p38/Hog1 MAPK and cell wall integrity (CWI) MAPK pathways and revealed that while the CWI MAPK pathway influences cell wall masking at the post-translational level, the p38/Hog1 pathway influences translatome reprogramming. We also identified another kinase, Yck2, that demonstrated robust β-1,3-glucan unmasking. We constructed a *YCK2* null mutant in our H99 background and *yck2*Δ:*YCK2* complemented strain and confirmed that Yck2 contributes to β-1,3-glucan masking at host temperature (**Figure 1 A, B**). The *yck2*Δ mutant demonstrated reduced growth compared to the wild type when grown at 37°C on solid media (**Figure 1C**). Kinetic growth assays demonstrate the *yck2*Δ mutant grows significantly slower than wildtype at 30°C and this growth defect is exacerbated at 37°C (**Figure 1 D, E**).

**Figure 1.**
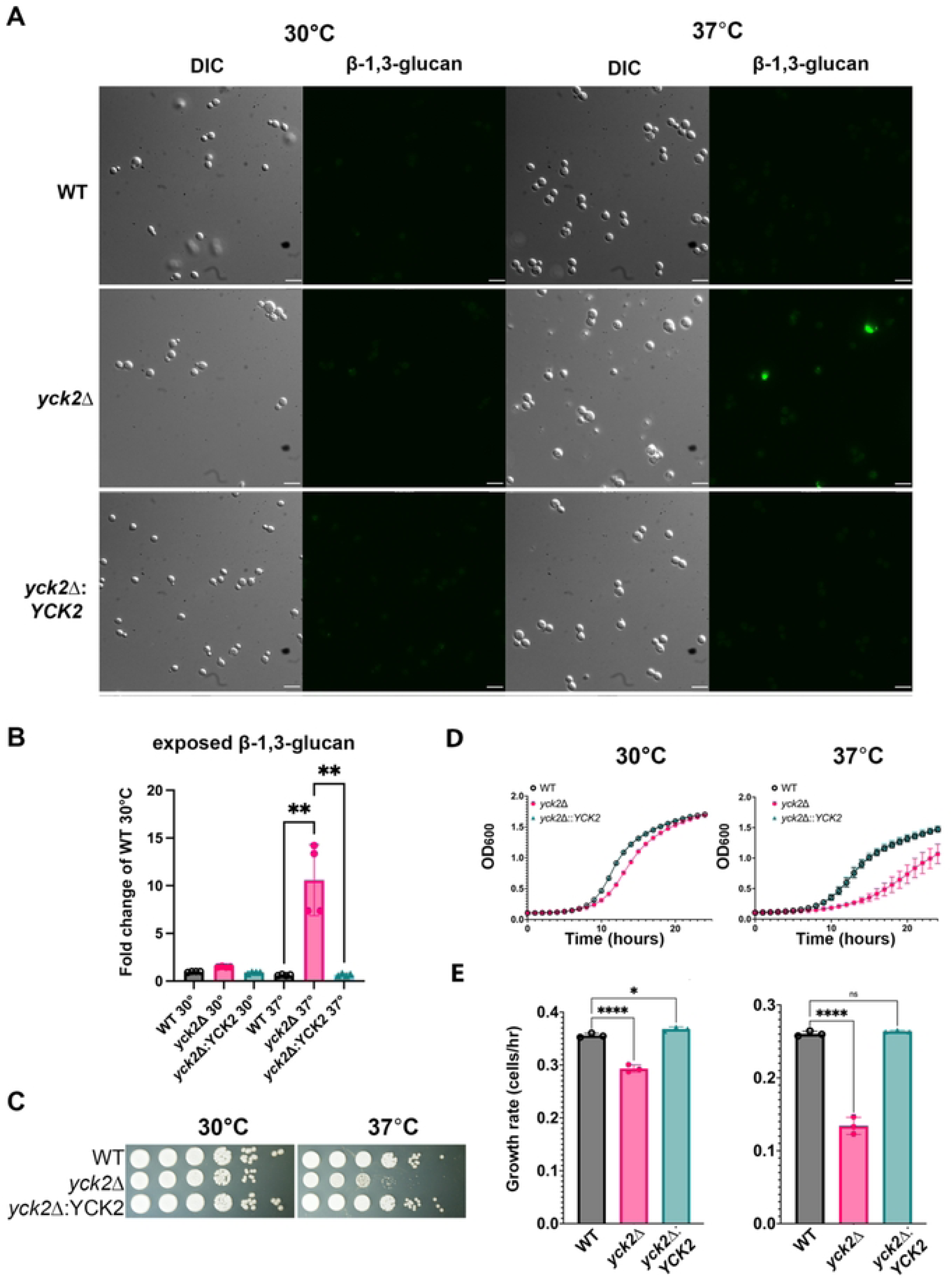
Absence of YCK2 results in β -1,3-glucan unmasking and reduced growth at host temperature. **A.** WT, *yck2*Δ, and *yck2*Δ:*YCK2* cells were grown to midlogarithmic phase, stained for β-1,3-glucan, and visualized by microscopy. Scale bar = 10um, images are representative of 3 biological replicates. **B.** Cells stained for β-1,3-glucan were assessed by flow cytometry. Statistical differences were determined by Kruskal-Wallis with Dunn’s test post hoc; *n* = 4, error bars = SD. **C.** Growth of cells at 30°C and 37°C was assessed by the spot dilution method. **D.** Kinetic growth of cells grown at 30°C and 37°C was assessed by continuous growth in a temperature regulated plate reader over 24 hours; *n* =3, error bars = SD. **E.** Differences in growth rate during the exponential phase (∼8-13 hrs) were determined as described in methods and significance was determined by Kruskal-Wallis with Dunn’s test post hoc; *n* = 3, error bars = SD.

To determine if Yck2 contributes to unmasking via a role in translational reprogramming we examined hallmark signatures of proper reprogramming following a shift from 30°C to 37°C: 1) repression of ribosomal protein transcripts, 2) translational repression via phosphorylation of the translation initiation factor eIF2α, and 3) a changed translational landscape skewed toward repression, indicated by increased 60S polysome peaks and decreased polysomes. None of these key contributing factors of translational reprogramming were different in the *yck2*Δ mutant compared to WT, indicating that Yck2 does not significantly contribute to these specific processes in response to host temperature (**Figure S1**) and that the glucan exposure in the *yck2*Δ mutant arises from Yck2 regulation independent of translatome reprogramming.

To investigate the mechanism underlying the glucan unmasking in the *yck2*Δ mutant, we further phenotypically assessed cell wall properties by the spot dilution method in the presence of cell wall inhibitors. The *yck2*Δ mutant was sensitive to SDS, calcofluor white and congo red (**Figure 2A**). Congo red binds β-1,3-glucan and calcofluor white binds chitin, and both are reported to inhibit chitin synthesis (22). We assessed cell wall chitin, chitosan, and exposed chitooligomers by staining cells with calcofluor white, eosin Y, and wheat germ agglutinin, respectively, and measured levels in each strain by flow cytometry. While it has been shown previously that host temperature increases chitin, we observed 2-fold higher levels in the change in chitin composition when the *yck2*Δ mutant was grown at 37°C compared to wild type and complemented strains (**Figure 2B, Figure S2**). We also observed 4-fold higher levels in the change in chitosan and exposed chitooligomers in the mutant (**Figures 2B, S2, S3**).

**Figure 2.**
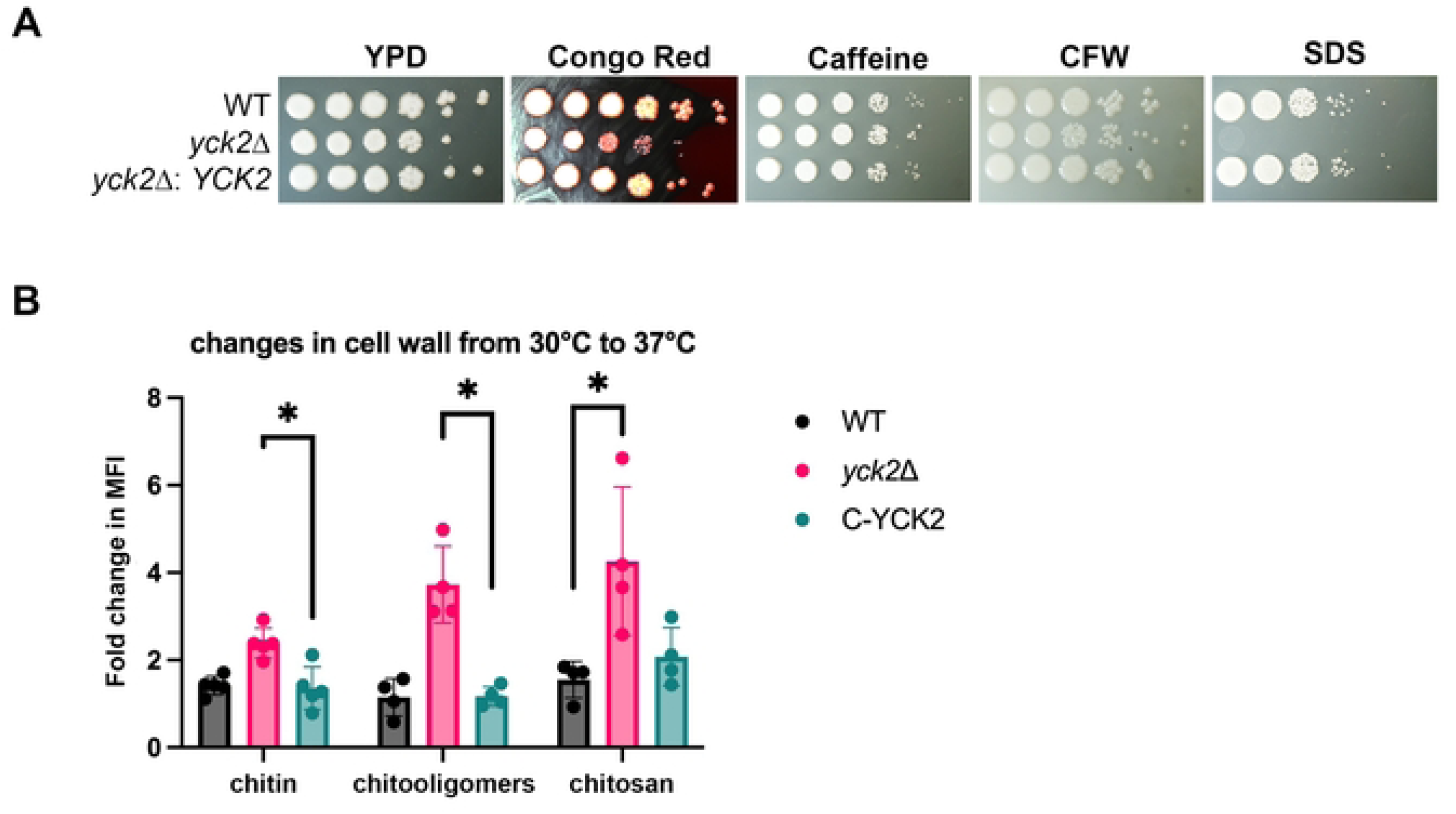
The *yck2*Δ mutant displays defective cell wall architecture. **A.** Cells were serially diluted and spotted onto YPD supplemented with 5 mg/ml congo red, 0.5 ml caffeine, 1 mg/ml calcofluor white (CFW), or 2% SDS. Plates were incubated at 30C for 3 days. Photographs are representative of 3 biological replicates. **B.** Cells grown to midlogarithmic phase at 30°C and 37°C were stained for calcofluor white, wheatgerm agglutinin, or eosin-Y, to detect levels of chitin, exposed chitooligomers, or chitosan, respectively. The fold change in MFI from cells grown at 30°C to 37°C is plotted and statistical differences were determined by Kruskal-Wallis with Dunn’s test post hoc; *n* = 4, error bars = SD.

### Yck2 regulates antifungal susceptibility

The antifungal drugs currently available target components of the fungal cell wall or cell membrane. Given the defect in cell wall integrity, we hypothesized that deletion of *YCK2* would sensitize *C. neoformans* to antifungal drugs. Growth on media containing the azole, fluconazole; the polyene, Amphotericin B; and the echinocandin, caspofungin, demonstrated that the *yck2*Δ mutant is sensitive to each of these classes of drugs (**Figure 3A**). The mutant was also sensitive to fludioxonil, an activator of *C. neoformans* Hog1 kinase and broad-spectrum fungicide used in agriculture. Notably, cryptococcosis cannot be treated with echinocandins due to the inherent resistance in *C. neoformans*; however, the *yck2*Δ mutant demonstrated sensitivity to caspofungin at concentrations 4-fold lower than WT suggesting that Yck2 is involved in processes that contribute to resistance.

**Figure 3.**
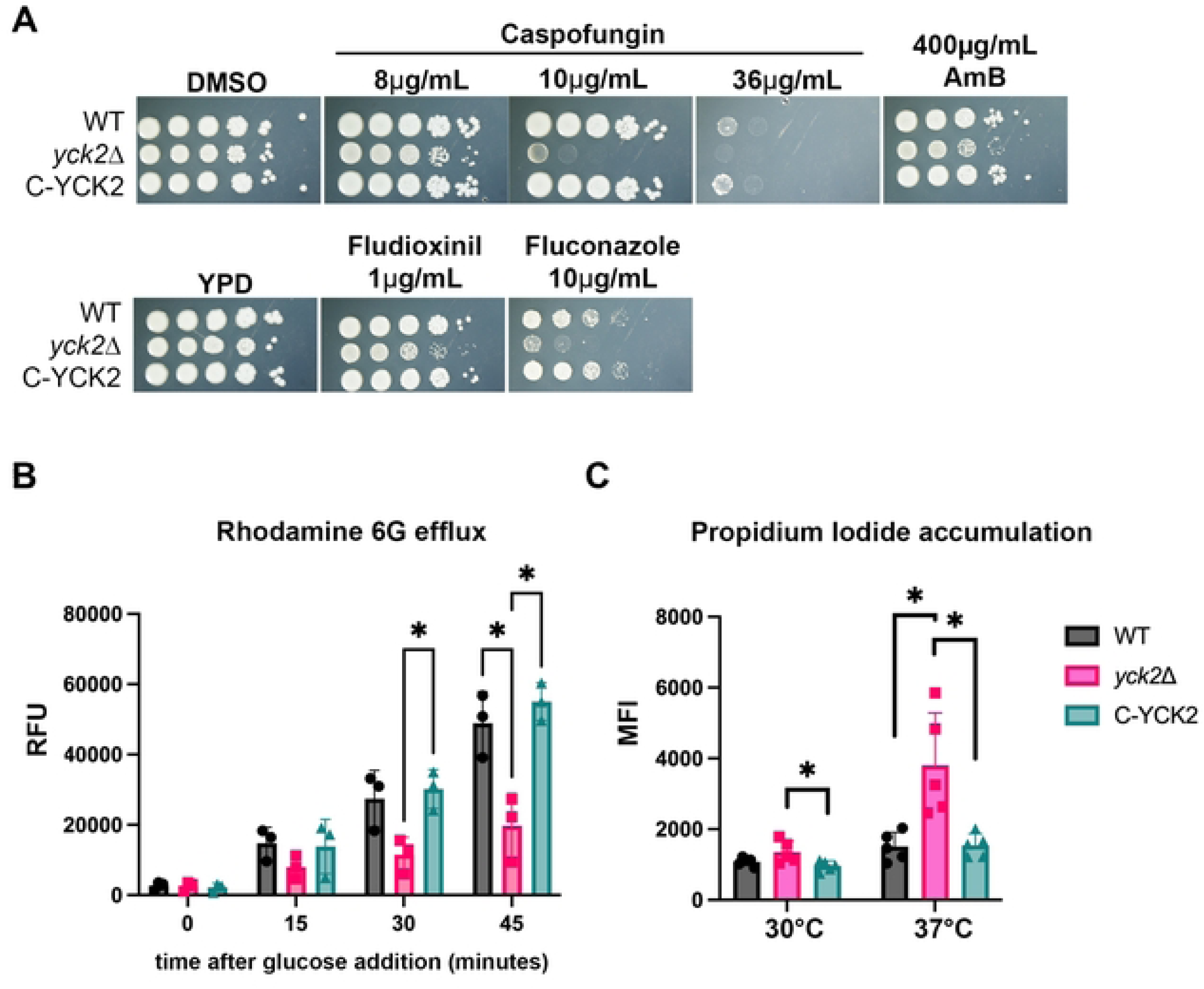
The *yck2*Δ strain exhibits increased antifungal susceptibility. **A.** Cells were serially diluted and spotted onto YPD agar supplemented with indicated antifungals and plates were incubated at 30°C for 3 days. Images are representative of 3 biological replicates. **B.** Starved cells were incubated with Rhodamine 6G (R6G) followed by supplementation of glucose. R6G efflux was measured by fluorescence of the supernatant. Statistical differences were determined by 2-way ANOVA with Tukey’s test post hoc; *n* = 3, error bars = SD. **C.** Permeability of cells was assessed by flow cytometric measurement of mean fluorescence intensity (MFI) of accumulated propidium iodide in cells grown at 30°C or 37°C to midlogarithmic phase. Statistical differences were determined by Kruskal Wallis with Dunn’s test post hoc; *n* = 5, error bars = SD.

Given the increased susceptibility of the *yck2*Δ mutant to all tested antifungals, we hypothesized that there may be a general defect in drug efflux in the *yck2*Δ mutant. To test efflux, we allowed starved cells to accumulate the fluorescent dye rhodamine-6-G (R6G), then added glucose and measured its efflux via the increase in R6G fluorescence in the supernatant. The *yck2*Δ mutant demonstrated significantly reduced R6G expulsion at each time point compared to wildtype and the complemented strains (**Figure 3B**). These results demonstrate that Yck2 promotes energy-dependent efflux in *C. neoformans* and may, in part, explain the antifungal sensitivity demonstrated by the *yck2*Δ mutant. We also tested whether Yck2 affects cell membrane permeability, which could also contribute to drug susceptibility. While there was no appreciable difference in permeability between the strains grown at 30°C, we observed a significant increase in permeability in the *yck2*Δ mutant compared to the wildtype and complemented strains when cells were grown at 37°C (**Figure 3C**).

### Yck2 regulates expression of genes involved in amino acid biosynthesis and redox

To begin to understand how Yck2 promotes cellular regulation we conducted RNA sequencing on wild type and the *yck2*Δ mutant grown under optimal conditions (30°C, YPD) and following a shift to host temperature for 1 hour considering the 37°C growth defect phenotypes and β-1,3-glucan exposure at host temperature. Lists of DEGs can be found in **Supporting Information S1**. While no terms were significant amongst upregulated differentially expressed genes (DEGs) in the *yck2*Δ mutant under optimal conditions, GO analysis demonstrated a significant reduction in genes involved in amino acid biosynthesis and oxidoreductase activity, both of which could imply defective mitochondria (**Figures 4A, 4B**). Notably, a study in *C. albicans* similarly found that Yck2 regulates amino acid biosynthesis and influences cell wall morphology suggesting conserved function (18). Surprisingly, GO analysis did not identify terms associated with cell wall biosynthesis.

**Figure 4.**
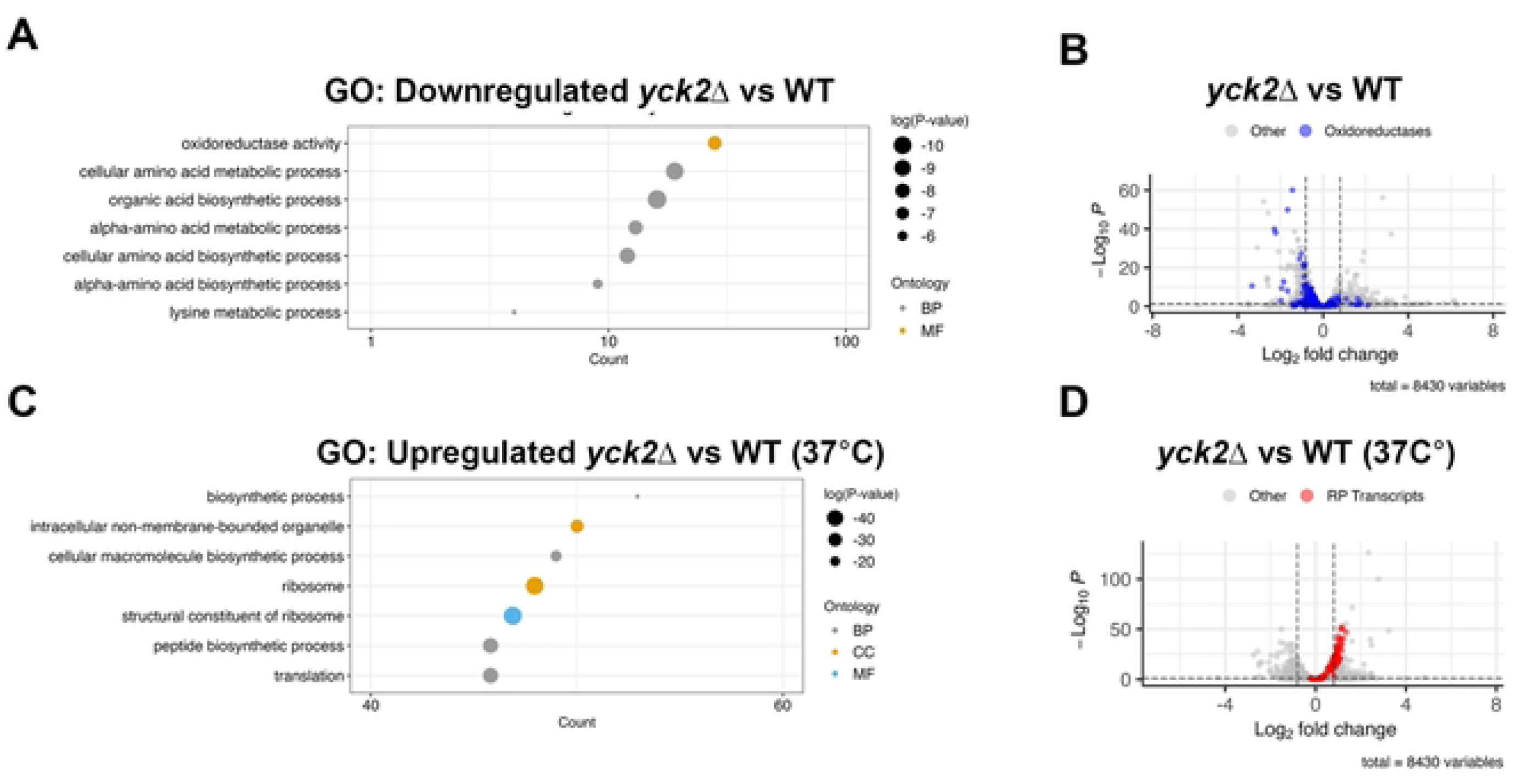
Yck2 regulates expression of genes involved in redox and amino acid biosynthesis. **A.** Gene ontology analysis of genes downregulated in the *yck2*Δ mutant compared to the wildtype strain when grown at 30°C. **B.** Volcano plot indicating genes with significant differences in expression in the *yck2*Δ strain compared to the wildtype strain. Vertical dashed lines indicate the 1.75 cut-off values. Blue circles indicate oxidoreductase genes. **C.** Gene ontology analysis of genes upregulated in the *yck2*Δ mutant compared to the wildtype strain when grown at 37°C. **D.** Volcano plot indicating genes with significant differences in expression in the yck2 strain compared to the wildtype strain when grown at 37°C. Vertical dashed lines indicate the 1.75 cut-off values. Red circles indicate genes encoding ribosomal proteins (RP).

Though not identified by GO analysis, we noticed that 2 genes encoding homologs of transporters involved in efflux in other fungi were downregulated in the *yck2*Δ mutant. CNAG_04576 is a homolog of the spermine transporter Tpo3, shown to promote azole resistance (23, 24) and CNAG_04098 encodes the homolog of Pdr15, belonging to the PDR family of ATP-binding cassette transporters that transport a broad range of chemicals out of cells (25, 26). We obtained mutants of *C. neoformans TPO3* and *PDR5* from *C. neoformans* deletion collections and examined sensitivity to antifungals to determine if their downregulation may contribute to the sensitivity seen in the *yck2*Δ mutant. Neither the *tpo3*Δ nor *pdr15*Δ mutant displayed more sensitivity to any antifungals compared to their parental strain (H99 for *pdr15*Δ and KN99 for *tpo3*Δ) suggesting that Yck2 influences efflux via means other than downregulation of these genes (**Supplementary Figure S4**).

GO analysis of downregulated DEGs in the mutant compared to WT following the temperature shift were similar to those observed in the unstressed condition, although additional downregulated oxidoreductases were identified. Upregulated DEGs in the mutant following the temperature shift were predominantly involved in ribosome biogenesis (**Figure 4 C, D**). We previously demonstrated that repression of abundant RP transcripts is important for translational reprogramming following host temperature stress (11, 27). As aforementioned, this response does occur in the *yck2*Δ mutant (**Supplementary Figures S1, S5**), however 60 minutes following the shift RP transcripts are more robustly decreased in the wild type than in the *yck2*Δ mutant which is reflected in the RNA-seq data. Overall, our RNAseq data suggest that Yck2 influences expression of genes involved in amino acid biosynthesis and cellular redox but does not appreciably regulate the rapid changes that occur in response to a shift to host temperature.

### Yck2 interacts with mitochondrial proteins

Since Yck2 is a kinase, we set out to identify potential targets of Yck2 using TurboID proximity-dependent biotinylation. We complemented the *yck2*Δ mutant with C-terminally myc+TurboID-tagged *YCK2* (**Figure 5A**) and grew this strain constitutively at 30°C or at 37°C with exogenous biotin. We identified streptavidin captured biotinylated proteins in WT cultures and the YCK2-turboID cultures by LC/MS/MS (**Supporting Information S1**). In cells grown at 30°C we identified 106 proteins that had > 2-fold peptide-spectrum matches (PSM) in the Yck2TurboID samples compared to WT samples (**Supporting Information S1**). Interestingly, GO analysis revealed that several of these proteins are involved in mitochondrial function including electron transport (**Figure 5C**) suggesting Yck2 may interact with these proteins prior to their mitochondrial import or that Yck2, in addition to its role at the plasma membrane, also localizes to the mitochondria. Interestingly, *S. cerevisiae* Yck1/2 have been shown to localize to the mitochondrial matrix and the outer mitochondrial membrane where they interact with the cysteine desulfurase Nsf1, and the mitochondrial import protein Tom22, respectively (28, 29).

**Figure 5.**
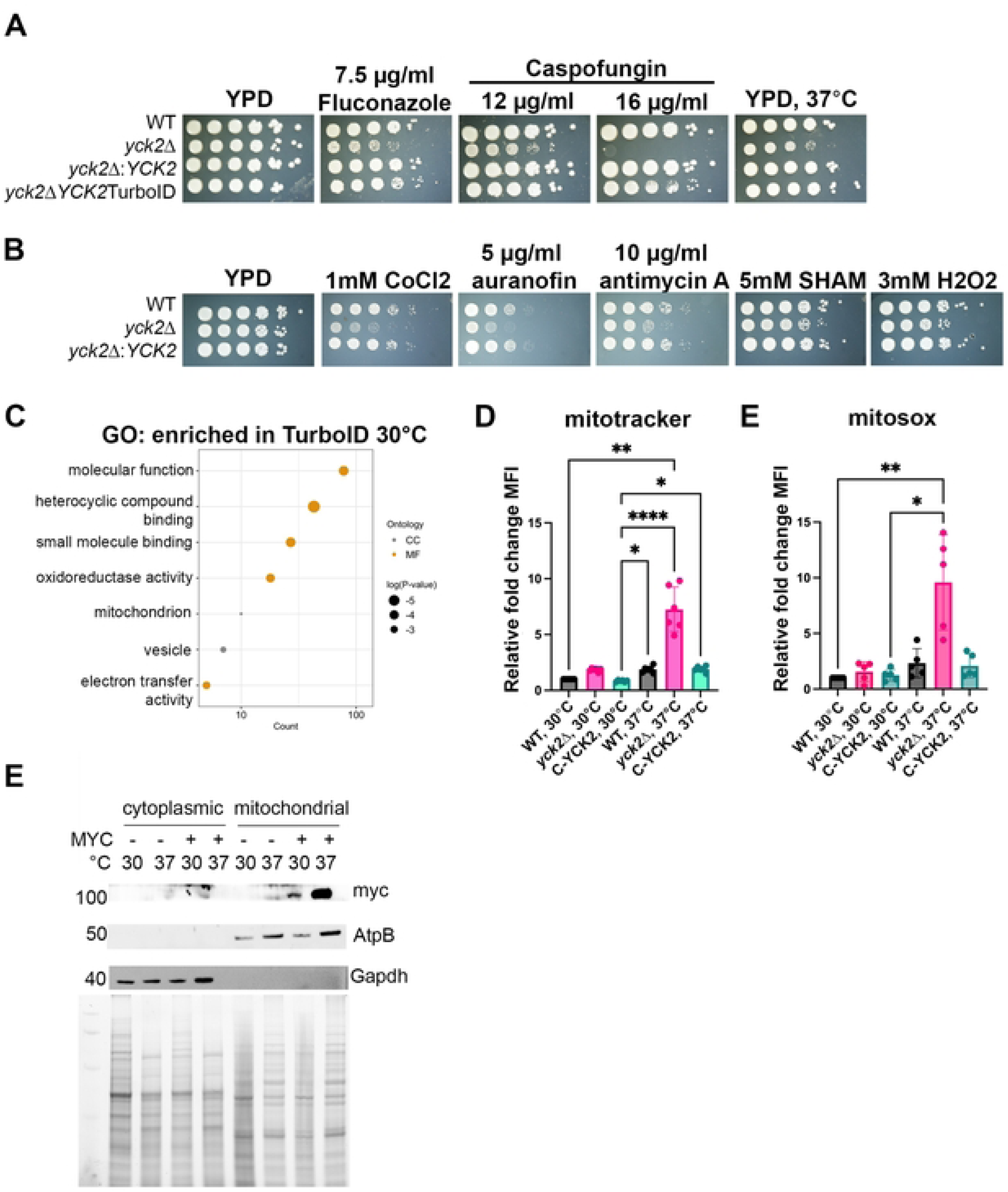
The Yck2 interactome suggests that Yck2 plays a role in mitochondrial regulation. **A.** TurboID-tagged Yck2 complements the *yck2*Δ mutant. Serially diluted cells were spotted and grown on YPD agar supplemented with indicated drugs and plates were grown at 30°C for 3 days. Images are representative of 3 biological replicates. **B.** Serially diluted cells were spotted and grown in YPD agar containing indicated mitochondrial inhibitors and plates were grown at 30°C for 3 days. Images are representative of 3 biological replicates. **C.** Gene ontology for proteins that were enriched in streptavidin pulldowns from Yck2TurboID cultures compared to wild type cultures. **D.** Cells grown to midlogarithmic phase at 30°C or 37°C were assessed for ROS production by Mitotracker CM-H2TMRos (Invitrogen) and analyzed by flow cytometry. Statistical differences were determined by Kruskal Wallis with Dunn’s test post hoc; *n* = 6, error bars = SD. E. Cells were grown and analyzed as in D but were stained for mtROS using mitoSox (Invitrogen). *n* = 5, error bars = SD. **F.** Western blots of cytoplasmic and mitochondrial subcellular fractions from the wild type and the *yck2*Δ:*YCK2*mycTurboID strain were probed for Yck2myc (myc), the mitochondrial marker AtpB, or the cytoplasmic marker Gapdh. For each sample 10μg of proteins was loaded as can be seen by the stain-free image under the blots.

### Yck2 regulates mitochondrial function and localizes to mitochondria

To gauge mitochondrial function, the wild type, *yck2*Δ, and *yck2*Δ:*YCK2* strains were grown in the presence of different mitochondrial inhibitors by the spot dilution method (**Figure 5B**). Antimycin A is an electron transport chain complex III inhibitor; CoCl_2_ is used to mimic hypoxia which interferes with oxidative phosphorylation; auranofin is an inhibitor of thioredoxin reductase, a major regulator of mitochondria redox balance; salicylhydroxamic acid (SHAM) is an inhibitor of the alternative oxidase, predominantly utilized under stress; and H_2_O_2_ is used to assess sensitivity to reactive oxygen species (ROS) which are predominantly produced in the mitochondria. The *yck2*Δ mutant demonstrated sensitivity to all these compounds suggesting that Yck2 plays a role in mitochondrial function. To specifically investigate if downregulation of oxidoreductases may impact cellular redox, we assessed levels of ROS with Mitotracker Cm-H_2_TMRos, a reduced derivative of dihydrotetramethyl rosamine, which is oxidized by cellular ROS causing fluorescence and is then sequestered in the mitochondria. When cells were grown at 37°C, but not 30°C, *yck2*Δ cells accumulated 4-fold higher levels of ROS compared to wild type suggesting that Yck2 is involved in regulating redox when grown at host temperature (**Figure 5D**). Most cellular ROS is derived from the mitochondria (30). Because Yck2 interacted with mitochondrially localized proteins and the mutant is sensitive to mitochondrial perturbation, we asked if Yck2 regulates mitochondrial ROS (mtROS) accumulation. We assessed mtROS using MitoSOX, which accumulates in mitochondria where it is then oxidized specifically by mitochondrially derived superoxide resulting in its fluorescence. Again, when cells were grown at 37°C, but not 30°C, *yck2*Δ cells accumulated 4-fold higher levels of mtROS compared to wild type (**Figure 5E**) suggesting that Yck2 specifically contributes to mitochondrial redox at host temperature.

To determine if Yck2 localizes to the mitochondria, we utilized the Yck2mycTurboID strain to determine if Yck2 is present in mitochondrial subcellular fractions. We were able to detect Yck2 in the mitochondrial fractions when grown at 30°C and 37°C, with greater accumulation at 37°C (**Figure 5F**). Yck2 was not detected in any of the cytoplasmic fractions likely due to its association with the plasma membrane.

### Yck2 interacts with Pma1 at 37°C and modulates intracellular pH

In cells grown at 37°C, the only unique hit from our TurboID proteomics was the plasma membrane H+ ATPase, Pma1, an essential gene that regulates membrane potential and cellular pH by pumping protons out of the cell. Complementing this finding, we noticed that amongst the upregulated genes in the mutant at 37°C were several involved in the Rim101 pH-responsive pathway (**Supporting Information S1)** suggesting that cellular pH may be dysregulated in the *yck2*Δ mutant. In *S. cerevisiae* Yck1/2 homologs phosphorylate Pma1 under glucose depleted conditions contributing to its inactivation. Interestingly, the Yck1/2 phosphorylation site on *S. cerevisiae* Pma1 located in the nucleotide-binding (N) domain is not conserved in *C. neoformans*, however the regulatory C-terminal region is extended in *C. neoformans* with several potential S/T residues that could be phosphorylated (**Figure S6 A, B**).

To determine if Pma1 is affected by loss of *YCK2* we created in-locus mNeonGreen tagged Pma1 strains in the wild type and *yck2*Δ backgrounds to examine levels and localization of Pma1. While Pma1 is localized to the plasma membrane, as shown previously (31, 32), in both strains when grown at 30°C or 37°C, levels of Pma1 are reduced in the *yck2*Δ background when cells are grown constitutively at host temperature (**Figures 6 A, B**) suggesting that Yck2 may regulate the abundance of Pma1.

**Figure 6.**
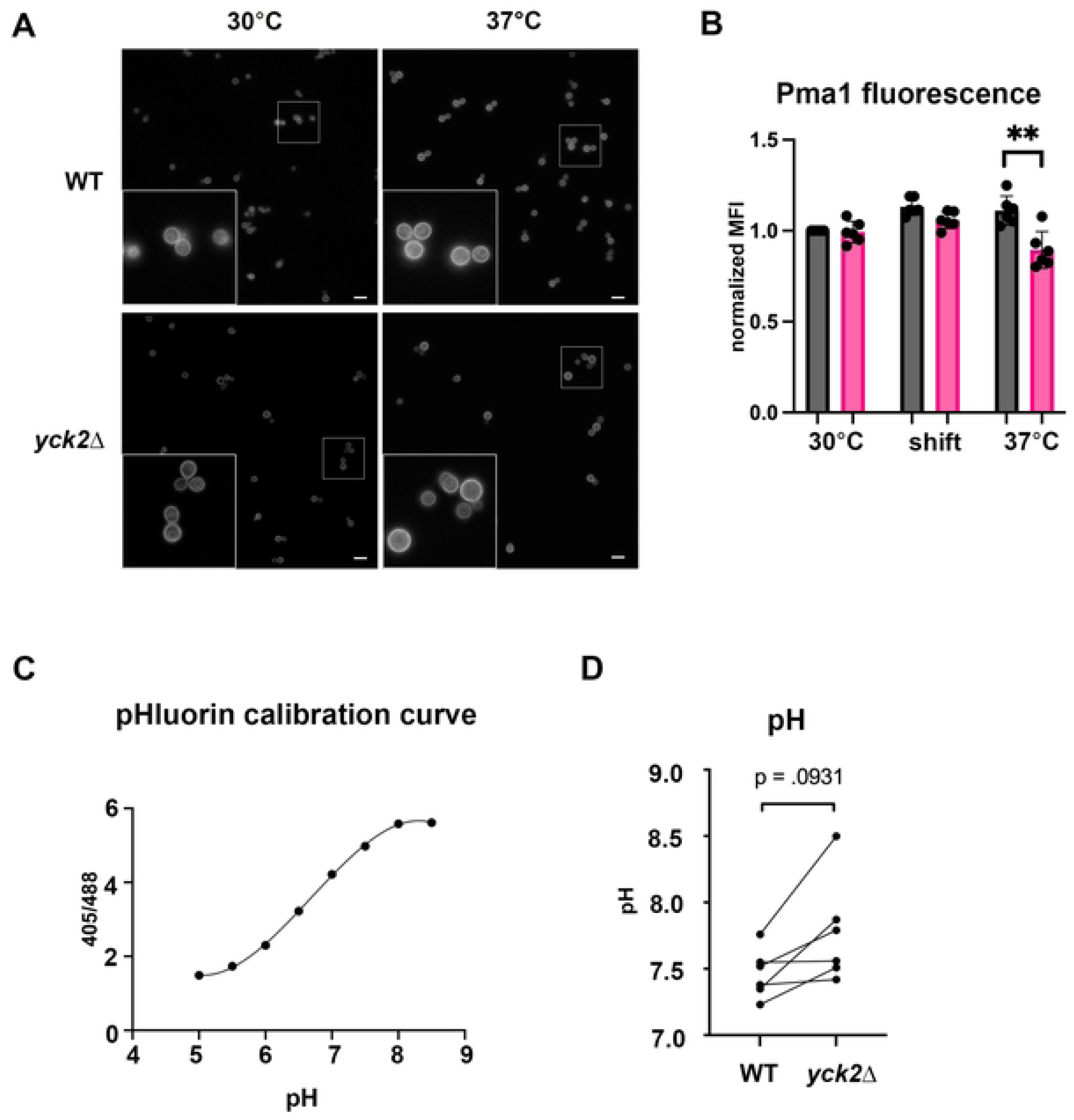
Yck2 regulates intracellular cellular pH. **A.** Pma1mNeonGreen-expressing WT and *yck2*Δ cells were assessed for Pma1 localization by fluorescence microscopy. **B.** Pma1mNeonGreen expressing cells were assessed for MFI by flow cytometry. Statistical differences were determined by Kruskal Wallis with Dunn’s test post hoc; n= 6, error bars = SD. **C.** Codon optimized pHluorin-expressing cells were subjected to incubation in buffers ranging from pH 5 - 8.5 to establish calibration curves determined by bimodal excitation at 405nm and 488nm and Em at 509nm. Image is representative of *n* = 6 biological replicates. **D**. pHluorin-expressing WT and *yck2*Δ cells grown in YPD at 37°C were assessed for intracellular pH by flow cytometry. The pH of each strain was converted to [H+] and the fold change [H+] in the mutant compared to WT was determined. Statistical difference was determined by one sample t-and Wilcoxen test; *n* = 6.

To assess if Yck2 affects the ability to regulate cytoplasmic pH, we created a strain expressing *C. neoformans* codon-optimized ratiometric pHluorin2 in our wild type background. Ratiometric pHluorin2 is a pH-sensitive derivative of GFP that has bimodal excitation at 395 and 475 nm and emission at 509 nm (33). Excitation at 395 nm decreases with a concomitant increase in excitation at 475 nm with increasing acidity allowing cellular changes in pH to be determined when using a standard curve achieved with ionophore-treated pHluorin-expressing cells incubated in buffers with known pH (**Figure 6C**). After verifying that our pHluorin2 strain responded to pH as previously described in other organisms we knocked out *YCK2* in the pHluorin2-expressing strain. We consistently observed higher intracellular pH in the *yck2*Δ:pHluorin2 strain compared to the pHluorin2 wild type strain (**Figure 6D**) suggesting that Yck2 prevents intracellular alkalinization by negatively regulating Pma1. We postulate that reduction of Pma1 protein levels that we observed in the mutant may be a response to help combat cellular alkalization.

### Loss of YCK2 does not impact early phagocytosis

A *yck2*Δ mutant strain (referred to as *cck1*Δ) demonstrated significantly attenuated virulence in an intranasal mouse model of infection (21). In this previous study, mice infected with *yck2*Δ did not begin to succumb to infection until after day 40, twice as long as mice infected with wild type. Given the changes in the cell wall we observed in the *yck2*Δ mutant, especially exposure of chitooligomers and β-1,3-glucan when grown at host temperature, we reasoned that the mutant would be more readily phagocytosed by macrophages. Our previous work demonstrated rapid uptake of the *ccr4*Δ mutant strain, which also exposes β-1,3-glucan, by alveolar macrophages compared to wild type just after 5 hours of infection (11). Surprisingly, when Pma1mNeonGreen-expressing wildtype and *yck2*Δ cells were incubated with THP-1 macrophages for 2 hours we did not observe statistical differences in uptake of *yck2*Δ cells compared to the wildtype (**Figure 7A**). We also examined uptake of cryptococcal cells by phagocytes present in the bronchoalveolar lavage (BAL) of BalbC/J mice infected intranasally with Pma1mNeonGreen or *yck2*Δ:Pma1mNeonGreen for 24 hours (**Supplementary Figure S7).** Though we observed a modest trend toward increased uptake of *yck2*Δ cells in macrophages and neutrophhils, there was no statistical difference in phagocytosis at these early timepoints (**Figure 7B**).

**Figure 7.**
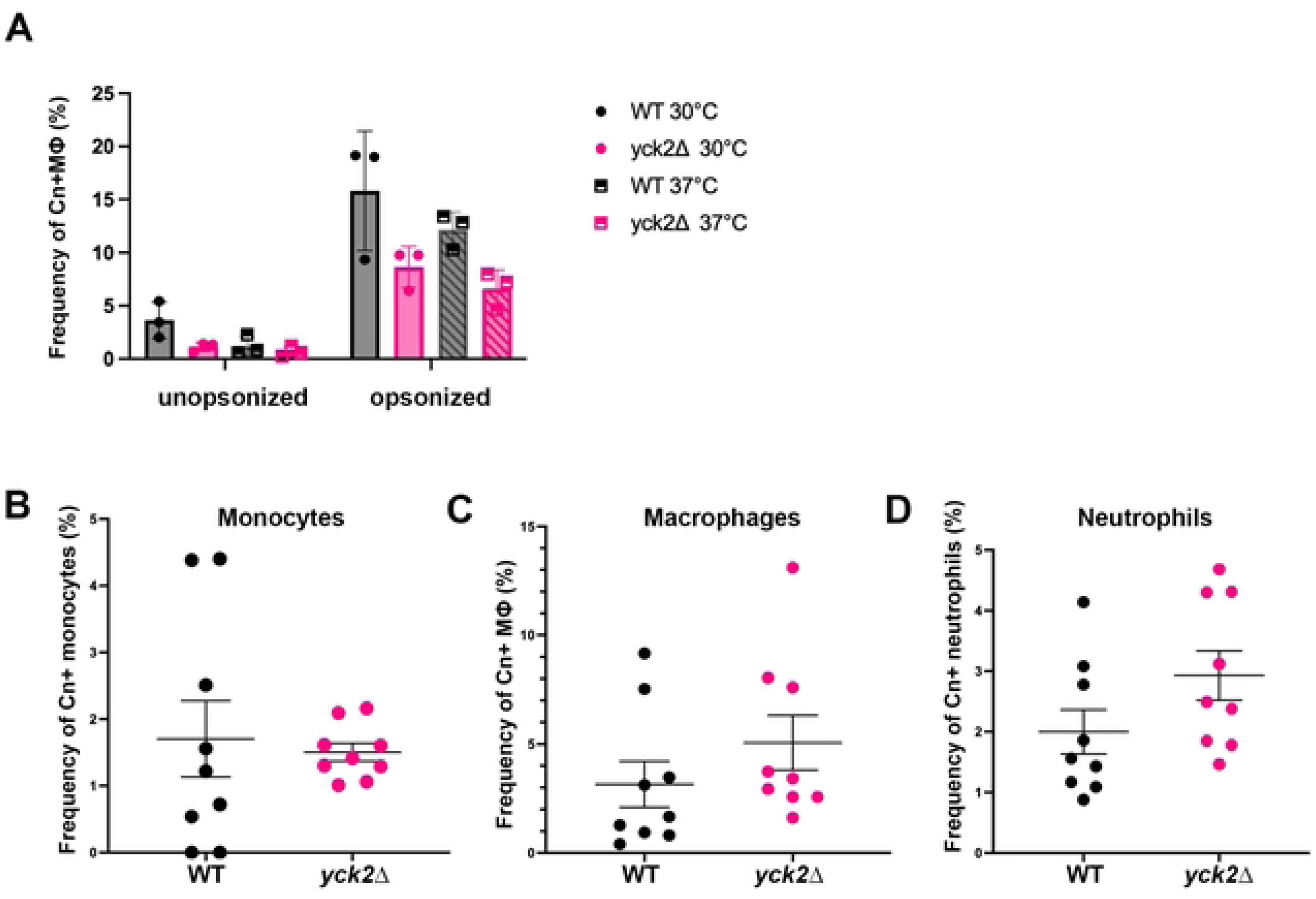
Loss of *YCK2* does not impact uptake by phagocytes at early timepoints. **A.** Opsonized or nonopsonized Pma1mNeonGreen and Pma1mNeonGreen:*yck2*Δ cells grown at 30°C or 37°C were incubated with THP-1 macrophages and uptake by macrophages was assessed by flow cytometry. *n* = 3. **B-D.** Uptake of cryptococcal cells in phagocytes in the BAL was assessed by flow cytometry. *n* = 9.

## Discussion

With growing limitations of the few current classes of antifungals, new therapeutic targets and regimens are imperative to control the rising rates of invasive fungal infections. Targeting kinases is an attractive antifungal avenue as they are hubs for cellular signaling. In this study we show that the casein kinase Yck2 contributes to a multitude of cellular processes in *C. neoformans* that may be due to downstream effects from its role in mitochondrial regulation and/or regulating intracellular pH. Work from the Cowen laboratory has recently revealed significant progress in derivatizing a small molecule Yck2 inhibitor identified from their previous drug screen (20). Their stabilized derivatives can be used in vivo and have high specificity to *C. albicans* Yck2 with limited effect on human CK1, demonstrating that fungal Yck2 is druggable. Yck2 homologs have pleiotropic effects in fungi that greatly influence virulence; thus, understanding the mechanisms by which Yck2 governs cellular processes will strengthen the efforts to develop and rationalize the use of Yck2-targeted antifungal therapies.

Our work supports roles for *C. neoformans* Yck2 in cell membrane integrity, antifungal resistance, cell wall architecture, and mitochondrial function. Several of these functions are temperature dependent as we see dysregulation in the *yck2*Δ mutant strain only when subjected to host temperature. Interestingly, the effect on antifungal drug susceptibility was not dependent on temperature. We hypothesize that Yck2 phosphorylation of unknown targets may alter drug uptake/efflux, or that the function(s) of Yck2 in the mitochondria, while not apparently substantial during growth at 30°C, may be critical in the context of triggers that elicit additional stress on the mitochondria, e.g. increased temperature or addition of drugs. Indeed, a complex relationship between mitochondrial function and antifungals has been established. In *Candida parapsilosis,* inhibition of mitochondrial function enhances caspofungin susceptibility (34), and in *C. albicans* electron transport chain complex I mutants display enhanced fluconazole sensitivity (35). Interestingly, in *Nakaseomyces glabratus* it was demonstrated that prior, but not simultaneous, mitochondrial inhibition reduces caspofungin tolerance suggesting that mitochondrial impairment could be a prerequisite for enhanced activity of some drugs in some fungi (36). In some instances, mitochondrial dysfunction leads to increased drug efflux and reduced antifungal susceptibility (37). While the *yck2*Δ mutant is deficient in energy-driven efflux from the cell it is unclear if Yck2 regulates the activity of transporters or contributes to the production of ATP needed to drive this efflux. Assessing cellular levels of ATP and performing phosphoproteomics under specific conditions in the future will help us further understand the mechanisms by which Yck2 regulates cellular processes including antifungal tolerance.

The cell wall and membrane ergosterol have historically been attractive targets for antifungal therapy due to the lack of homologous human components. Inhibition of Yck2, which increases antifungal susceptibility, may potentiate current antifungals and as our data suggests, could possibly enable the use of echinocandins in treatment of cryptococcosis. With the progression of fungal specific Yck2 inhibitors, as demonstrated in *C. albicans* (20), testing for potentiation in *C. neoformans* is a realistic and sensible endeavor.

We demonstrated that *C. neoformans* Yck2 is required for cell wall masking and architecture when cells are grown at host temperature. Specifically, levels of chitin, chitosan, exposed chitooligomers, and exposed β-1,3-glucan are elevated in the mutant. Like antifungal susceptibility, it is possible that mitochondrial dysfunction leads to the cell wall defects that we observed in the *yck2*Δ mutant as it is becoming increasingly apparent that defective mitochondrial function in fungi impacts cell wall morphology (38). For example, mutants in *C. albicans* lacking mitochondrial-localized proteins involved in electron transport chain display altered cell walls (39). Interestingly, the *C. neoformans* mitochondrial-localized protein Mar1, was shown to regulate ROS generation and cell wall morphology when grown in host-like conditions, similar to *yck2*Δ (40).

Unmasked pathogen-associated molecular patterns (PAMPs) can lead to recognition, clearance, and immune responses that dampen fungal virulence. Previous assessment of *yck2*Δ (*cck1*Δ) for virulence in a mouse model demonstrated that mice infected with *yck2*Δ survived more than twice as long as mice infected with wildtype before beginning to succumb to infection (21) which suggested to us that early responses are likely diverse. We previously demonstrated that a *ccr4*Δ mutant, which exposes β-1,3-glucan and mannoproteins (11, 41), was significantly more phagocytosed than WT in a mouse model after 5 hours; we thus reasoned that exposed β-1,3-glucan and chitooligomers, drivers of Dectin-1 and TLR2 signaling, respectively, would lead to increased phagocytosis of the *yck2*Δ mutant. However, we observed no difference compared to uptake of the wildtype using ThP-1 macrophages or assessing macrophages from the BAL of infected mice suggesting that PAMP exposure alone does not govern uptake at early timepoints. We note that in our previous study the exterior surface of cells were fluorescently labeled to allow detection, which could possibly have enhanced recognition and uptake.

Interestingly, the *C. neoformans mar1*Δ mutant is severely attenuated in virulence, and despite significantly reduced fungal burden in organs and cell wall defects of the *mar1*Δ mutant, the early pulmonary immune response of mice infected with *mar1*Δ was similar to mice infected with wildtype (42). This is strikingly similar to our results with the *yck2*Δ mutant. Authors found that the *mar1*Δ mutant however was found to have a predilection to cause granulomatous lung infection (40, 42, 43). Thus, assessing fungal burden, histology, and the immune response at later time points are necessary to better understand the effect of Yck2 on host-pathogen interactions. It will be interesting to assess if the *yck2*Δ mutant and other mutants of genes involved in mitochondrial function that also affect cell wall remodeling develop granulomatous lung infections similar to the *mar1*Δ mutant.

Our TurboID experiments also suggest that Yck2 interacts with the essential membrane H+ ATPase, Pma1. This was not surprising as Yck2 has been shown to regulate Pma1 in *S. cerevisiae*, however we revealed that the Yck1/2 site of phosphorylation on Pma1 is not conserved in *C. neoformans*, suggesting that Yck2 may phosphorylate Pma1 elsewhere, perhaps in the C-terminal regulatory domain. Pma1 is the major regulator of plasma membrane potential and pH homeostasis. Absence of *YCK2* did not impact localization of Pma1, but we did observe reduced levels in the mutant when cells were grown at host temperature. Interestingly, despite this reduction in Pma1 levels, we observed more alkaline intracellular pH in the mutant suggesting that, like in *S. cerevisiae*, Yck2 phosphorylation of Pma1 limits H+ pumping activity. Reduced levels of Pma1 in the mutant may be a means for the cell to combat its overactivity. Alternatively, higher activity via a Yck2-independent mechanism could be a means to compensate for Yck2-dependent reduced levels in the mutant. It is unclear if mitochondrial dysfunction is a downstream effect of alkaline intracellular pH signaling or if Yck2 has a more direct effect on the mitochondria; our interactome data suggests the latter. However, alkalinization of the cytosolic pH has been shown to dissipate the pH gradient in the mitochondria, resulting in aberrant oxidative phosphorylation and mitochondrial dysfunction (44). Future studies will investigate the localization of Yck2 in response to host like stress and identify phosphosites on Pma1 and other potential alternative targets to better elucidate the direct impact(s) of Yck2.

With rapidly increasing rates of invasive fungal infections and development of antifungal resistance along with the slow development of novel antifungal drugs it is critical to invest in understanding worthwhile targets such as Yck2. Yck2 is a versatile kinase involved in a multitude of critical cellular functions that aid in the ability of pathogenic fungi to adapt to stress including cell wall morphology, antifungal susceptibility and virulence. Now with a link to regulating cellular pH and mitochondrial function, we are beginning to understand the pleiotropism of this gene. The mitochondria are vital to most fungal pathogens and regulate adaptation to the host; thus, targeting the mitochondria may be a fruitful antifungal strategy. The impact of Yck2 in the mitochondria may create downstream effects that ultimately affect the rest of these processes, but additional work is needed to determine the interconnectedness of Yck2 functions. With advances being made for development of fungal specific Yck2 inhibitors, a focus on understanding how this kinase is orchestrating these processes is significant. Further, regulation of mitochondria is an attractive approach to antifungal therapy, and mitochondrial perturbation can alter the efficacy of current antifungals. Determining the mechanism by which *YCK2* deletion sensitizes *C. neoformans* to antifungal drugs will be significant for the development of combination therapy.

## Methods

### Strains and growth conditions

All strains generated in this study were constructed in the H99 wild type background. The *tpo3*Δ and *pdr5*Δ strains were obtained from the Madhani (45) and Lodge knock out collections, respectively, purchased from FGSC. Strains purchased from the FGSC were validated by PCR through the genomic loci. Strains were streaked from glycerol stocks onto YPD agar and grown at 30°C for 2-3 days. For culturing: a colony was resuspended in 4 mLs of YPD and grown at 30°C, 250 rpm overnight; overnight cultures grown in YPD were used to seed YPD or YNB+2% dextrose at OD_600_ = 0.2 and cells were grown to mid-logarithmic phase (OD_600_ = 0.6) at 30°C or 37°C, 250 rpm unless otherwise specified.

All oligonucleotide primers used in this study to create and verify strains can be found in Supporting Information S1.

The *yck2*Δ mutant was constructed by PCR amplifying the knockout construct from the *yck2*Δ strain from the Bahn kinase knock out collection (46) purchased from the FGSC with primers F-yck2KO-amp and R-yck2KO-amp. Knockouts were confirmed by PCR amplification of the genomic locus with primers F-YCK2KOscreen and R-YCK2KOscreen, and by northern and southern blot analyses. To complement the *yck2*Δ mutant *YCK2* with 1kb upstream and downstream sequence was PCR amplified from H99 genomic DNA with primer F-YCK2-1kbUP-SpeI and F-YCK2-1kbDOWN-SpeI and ligated into plasmid pBluescript + NEO. The plasmid was transformed into the mutant by biolistic transformation (47) and complementation was verified by northern blot and restoration of phenotypes via the spot dilution method.

The Yck2-turboID construct was created using the NEBuilder system. Yck2 was amplified with F-YCK21kbUP-SpeI and R-YCK2 NEBuild; the TurboIDmyc sequence from pFB1434 (Addgene plasmid no. 126050) (48), was amplified with primers F/R-TurboID-NEBuild; the Yck2 terminator was amplified with primers F-YCK2term-NEBuild and R-YCK2term-SpeI. The product was digested with SpeI and ligated into pBluescript+NEO for biolistic transformation into the *yck2*Δ mutant. Complementation was verified by northern blot, western blot, and phenotypic assessment by spot dilution analyses.

To create the pHluorin2 expressing strain, the pHluorin2 sequence (49) was first codon-optimized for *C. neoformans* and an intron from CNAG_05429 was placed within the sequence. *C. neoformans* optimized pHluorin2 was then synthesized in pUC57 with the *ACT1* promoter (p) and *GPD1* terminator (t) by Genscript. The *p*ACT-pHluorin-GPD1*t* construct was PCR amplified from the pUC57+pHluorin plasmid, inserted into pBluescript+NAT and transformed into H99 by biolistic transformation. To make a pHluorin:*yck2*Δ strain ∼500 bp upstream and downstream of *YCK2* were PCR amplified from H99 genomic DNA using primers listed in Table SX. The G418 resistance cassette was PCR amplified from the pBluescript+NEO plasmid. PCR products were digested, purified, and ligated into pBluescript. The full knock-out construct was PCR amplified and used to transform the pHluorin expressing strain. Knock out of *YCK2* was verified by PCR amplification through the locus and northern blot.

The Pma1mNeonGreen strains were constructed by CRISPR as described previously (32) with the exception that we used a modified CRISPR_sg/R oligonucleotide (Supporting Information S1).

### Spot dilution assays

Overnight cultures grown for 16-18 hours were pelleted, washed twice with sterile deionized water (SDW), and resuspended in SDW to OD_600_ = 1.0. Six 10-fold serial dilutions were made in sterile water and 5 were spotted onto agar plates supplemented with indicated drugs and incubated for 3 days.

### Growth curves

Overnight YPD cultures were diluted to OD_600_ = 0.1 in YPD and 100μL was dispensed into 96-well plates. Plates were incubated in a Biotek Synergy plate reader with continuous linear shaking at 30°C or 37°C and OD_600_ was recorded every 15 minutes for 24 hours. For each biological replicate 3 technical replicates were plated and averaged. Growth rate during exponential growth (∼8–13h) was calculated across three biological replicates as follows: Rate = ln(OD_600_(final)/ OD_600_(initial))/duration.

### Cell staining, flow cytometry, and microscopy

For cell staining of chitin and WGA, 500 μL of mid-logarithmic phase cultures grown in YPD were pelleted and washed twice with PBS. Cells were fixed with 3.7% formaldehyde for 5 min, washed twice with PBS, and resuspended in 15 μg/ml WGA in PBS. Cells were incubated in the dark for 15 min followed by the addition of calcofluor white at 25 μg/ml and 10 min of incubation in the dark. Cells were washed twice with PBS and resuspended in PBS. For staining cell wall chitosan, 500 μL of mid-logarithmic phase cultures were pelleted and washed three times with McIlvaine’s buffer (pH 6), resuspended in 150 ug/ml Eosin Y in McIlvaine’s Buffer, and incubated in the dark for 10 min. Cells were washed and resuspended in McIlvaine’s buffer.

For staining exposed β-1,3-glucan, 500 μL of mid-logarithmic phase cultures grown in YPD were pelleted and washed twice with PBS. Cells were resuspended in 15 ug/ml anti-β-1,3-glucan antibody (Biosupplies Australia) or 15 ug/ml IgG-kappa isotype control and incubated in the dark with agitation for 45 min. Cells were washed three times with PBS, resuspended in 10 ug/ml Alexafluor 488 conjugated anti-mouse secondary antibody and incubated in the dark with shaking for for 30 min. Cells were washed twice with PBS, fixed with 3.7% formaldehyde, washed and resuspended in PBS.

For detection of ROS, 3 mLs of mid-logarithmic phase culture grown in YNB+2% dextrose was treated with 500 nM Mitotracker Orange CM-H_2_TMRos (Invitrogen) for cellular ROS, 5 μM MitoSOX (invitrogen) for mitochondrial ROS, or 200 nM Mitotracker Green FM (Invitrogen) for 30 minutes. Cells were washed twice and resuspended in PBS.

Flow cytometry was performed on a BD Fortessa flow cytometer. Analyses were performed using FlowJo v10. Fluorescence microscopy was performed using a Leica DM 6B upright fluorescence microscope.

### Northern blot

Midlogarithic cultures grown at 30°C were pelleted and resuspended in prewarmed 37°C YPD and incubated at 37°C, 250 rpm. 5 ml of culture was pelleted and flash frozen every 30 minutes for 2 hours and stored at -80°C until lysis. For RNA extraction, 50 μL of buffer RLT (Qiagen) + 10 uL/mL β-mercaptoethanol was added to cell pellets and cells were lysed by mechanical disruption with glass beads in a Bullet Blender Gold for 5 min. 250 μL of RLT + 10 uL/mL β-mercaptoethanol were added to lysed cells and the supernatant was clarified by centrifugation at 13,000 rpm for 2 min. Equivalent amounts of 70% ethanol was added to the clarified lysate and RNA was extracted using Qiagen RNeasy kit per manufacturer’s protocol. For northern blot, 3 μg of RNA was loaded onto a 1% agarose + formaldehyde gel supplemented with Sybersafe and RNA was electrophoretically separated at 80V for 30 min. The gel was imaged for rRNA detection using Biorad GelDoc XR. RNA was transferred to nylon and hybridized with a P32-labeled *RPL2* DNA probe. Blots were exposed to a phosphor screen and imaged using Licor Typhoon Imaging system.

### Polysome profiling

Cells were grown to mid-log phase in YPD at 30°C, and half of the culture was pelleted, resuspended in prewarmed 37°C YPD, and incubated at 37°C for 60 min. Pellets were flash frozen in liquid nitrogen and stored at −80°C until lysis. Pellets were washed in polysome lysis buffer (20 mM Tris HCl, pH 8.0, 140 mM KCl, 5 mM MgCl_2_, 1% Triton X-100, 25 mg/ml heparin sodium sulfate, 0.1 mg/ml cycloheximide) and pelleted. Cells were resuspended in residual buffer, transferred to a microcentrifuge tube, and pelleted at 13,000 rpm for 30 s. The supernatant was removed, and pellets were resuspended in 100 μl of cold lysis buffer and transferred to an Eppendorf tube containing glass beads. Cells were lysed mechanically in a Bullet Blender for 5 min, followed by the addition of 300 μl cold lysis buffer. The supernatant was transferred to a cold microcentrifuge tube and centrifuged for 10 min at 14,000 rpm and 4°C. Cleared lysates were quantitated for RNA, and 250 μg was loaded on top of 10% to 40% sucrose gradients (in 20 mM Tris HCl, pH 8.0, 140 mM KCl, 5 mM MgCl_2_, 25 mg/ml heparin sodium sulfate, 0.1 mg/ml cycloheximide). Gradients were subjected to ultracentrifugation for 2 h at 39,000 rpm and 4°C. Sucrose gradients were then run through a Piston Gradient Fractionator (Biocomp) measuring *A*_260_.

### Western blot

Midlogarithic cultures grown at 30°C in YPD were pelleted and resuspended in prewarmed 37°C YPD and incubated at 37°C, 250 rpm. 15 ml of culture was pelleted and flash frozen every 15 minutes for 60 min and stored at -80°C until lysis. 50 μL of protein extraction buffer (15 mM HEPES [pH 7.4], 10 mM KCl, 5 mM MgCl_2_, 10 μl/ml Halt protease inhibitor, 1 mM dithiothreitol [DTT]) was added to pellets and cells were lysed by mechanical disruption with glass beads in a Bullet Blender Gold for 5 min. 100 μL of lysis buffer was added to lysed cells and the supernatant was centrifuged for 10 min at 13000 x g, 4°C. Clarified lysate was quantified and 10 μg of protein was electrophoretically separated through a stain-free polyacrylamide gel (Biorad). Total protein was quantified by activation of stain-free reagent and imaged with Biorad Gel Doc. Protein was transferred to nitrocellulose using the Biorad trans-blot Turbo system, blocked with Everyblot Blocking Buffer (Biorad), probed with Rabbit anti-eIF2α antibody ((Genscript) 1:1,000, 4°C, overnight) followed by HRP-conjugated anti-Rabbit IgG, and imaged by chemiluminescence on Biorad ChemiDoc. The blot was stripped and probed with Rabbit anti-Phosphorylated-eIF2α antibody ((Abcam) 1:1,000, 4°C, overnight), followed by HRP-conjugated anti-Rabbit IgG, and imaged as above.

### Rhodamine 6G Efflux Assay

Cells were grown in YPD to mid-logarithmic phase, washed 3 times with sterile deionized water, and resuspended in PBS at room temperature for 2 hours. Rhodamine 6G was added to a final concentration of 10 uM and cells were incubated at 30°C, 250 rpm for 30 min. Cells were washed 3 times with PBS, resuspended in PBS and split into two tubes. Glucose was added to one tube at a final concentration of 2%. 250 μL of culture was pelleted every 15 minutes and the supernatant was placed on ice. 100 μL of supernatant for each time point (2 technical replicates) was transferred to a Black bottom 96-well plate and analyzed for fluorescence (Ex 525nm/Em 555nm) in a Biotek Synergy plate reader.

### Affinity purification of biotinylated proteins by TurboID-tagged Yck2

H99 and *yck2*Δ:YCK2TurboID-3xMyc cells were grown for 18-20 hours (OD_600_ = 1.5-1.8) at 30°C or 37°C, shaking (200 rpm) in 2 L YPD medium supplemented with 50 µM biotin. Cells were pelleted and flash frozen at -80°C. Frozen cell pellets were dislodged with 2 mL of RIPA buffer (50 mM Tris-HCl [pH 7.5], 150 mM NaCl, 1.5 mM MgCl_2_, 1 mM EGTA, 0.1% SDS, 1% NP-40, 0.4% sodium deoxycholate, 1 mM dithiothreitol (DTT), 1 mM phenylmethylsulfonyl fluoride, 1× cOmplete (Roche), and 1× PhosSTOP (Roche)) and lysed using a Krups coffee grinder for 2 min followed by grinding with mortar and pestle for 20 min. Liquid nitrogen was added as necessary to keep the lysate cold. Ground cell powder was transferred to a 50 mL conical, resuspended in 10 mL RIPA buffer, and incubated at 4°C for 20 minutes with end-to-end rotation. Lysates were treated with 750 U Benzonase to digest RNA and DNA for 1 hour. Clarified lysates were then incubated with 100 µL streptavidin-Sepharose for 4 hr at 4°C. Beads were washed four times with cold RIPA buffer followed by four washes with 20 mM ammonium bicarbonate. Beads were resuspended in ammonium bicarbonate and stored at −80°C until liquid chromatography tandem mass spectrometry (LC-MS/MS) analysis.

### LC-MS analysis

The LC-MS system consists of a Dionex Ultimate 3000 nano-LC system, a Dinex Ultimate 3000 micro-LC system with a WPS-3000 autosampler, and an Orbitrap Fusion Lumos mass spectrometer. A large-inner diameter (i.d.) trapping column (300-μm i.d. by 5 mm) was implemented before the separation column (75-μm i.d. by 65 cm, packed with 2.5-μm Xselect CSH C18 material) for high-capacity sample loading, cleanup, and delivery. For each sample, 8 μL derived peptides was injected for LC-MS analysis. Mobile phases A and B were 0.1% FA in 2% ACN and 0.1% FA in 88% ACN. The 180-min LC gradient profile was 4% for 3 min, 4–11% for 5 min, 11–32% B for 117 min, 32–50% B for 10 min, 50–97% B for 5 min, and 97% B for 7 min and then equilibrated to 4% for 27 min. The mass spectrometer was operated under data-dependent acquisition mode with a maximal duty cycle of 3 s. MS1 spectra was acquired by Orbitrap under 120-k resolution for ions within the *m/z* range of 400–1,500. Automatic gain control and maximal injection time were set at 120% and 50 ms, respectively, and the dynamic exclusion was set at 45 s and ±10 ppm. Precursor ions were isolated by quadrupole using a *m/z* window of 1.2 Th and were fragmented by high-energy collision dissociation for back-to-back Orbitrap/ion trap MS2 acquisition. Orbitrap MS2 spectra were acquired under 15-k resolution with a maximal injection time of 50 ms, and ion trap MS spectra was acquired under rapid scan rate with a maximal injection time of 35 ms. Detailed LC-MS settings and relevant information can be found in a previous publication by Shen et al (50).

### Mitochondrial Fractionation

Mitochondria were isolated as previously described (51) with some modifications. Cells were grown in YPD at 30°C or 37°C to midlogarithmic phase, flash frozen and stored at -80°C in 30% glycerol. Cells were washed twice in PBS, resuspended in 2 ml/g wet weight of DTT buffer (100 mM Tris [pH 7.4], 10 mM DTT) and incubated at 30°C for 20 minutes 70 rpm. Cells were pelleted and washed in 6.5 ml/g wet weight of SCS buffer (100 mM Sodium citrate [pH 5.5], 1.1 M sorbitol), resuspended in 6.5 ml/g wet weight filter sterilized lysis buffer (SCS buffer supplemented with 64 mg/ml VinoTaste (Bucher-Vaslin), 1 mg/ml Driselase (Medchem Express), 1 mg/ml β-glucanase from Trichoderma (Sigma) and incubated at 30°C, 70 rpm, for 3 hours to generate spheroplasts. Spheroplasts were pelleted and washed in 7 ml/g wet weight ice cold homogenization buffer (10 mM Tris [pH, 7.4], 0.6 M sorbitol), resuspended in homogenization buffer supplemented with 10ul/ml HALT protease and phosphatase inhibitor (Pierce), lysed by 20 strokes of dounce homogenization, then diluted 1:1 with cold homogenization buffer supplemented with HALT. Cellular debris, cytoplasmic fractions, and mitochondrial fractions were obtained by stepwise centrifugation as described. For protein detection, 10 μg of protein was electrophoretically separated in a 10% TGX Stain-free polyacrylamide gel (Biorad) and transferred to nitrocellulose. For detection of Yck2mycTurbo, mitochondrial AtpB,and Gapdh, membranes were probed with monoclonal anti-MYC antibody (Millipore, 1:1000 Biorad Everyblot Blocking buffer), polyclonal anti-AtpB antibody (Agrisera, 1:25,000 Biorad Everyblot Blocking buffer), or polyclonal anti-Gapdh (Abcam, 1:2500 3% BSA), respectively. Membranes were probed with HRP-conjugated secondary antibodies and imaged by chemiluminescence using a Biorad chemidocMP.

### Protein modeling and phosphosite determination

PDB files of the *S. cerevisiae* Pma1 protein and *C. neoformans* Pma1 protein, obtained from RCSD PDB and alphafold, respectively, were uploaded to PyMol (version 3.0.3). CnPma1 was aligned to ScPma1. The N-domain region bearing the Yck1/2 phosphosite in *S.cerevisiae* was used to locate the N-domain for *C. neoformans* Pma1. The 2 sequences were then aligned with Expasy alignment tool.

### Intracellular pH determination by pHluorin fluorescence

Measurement of pH using pHluorin was assessed by adapting the method described by Triandafillou and Drummond (52). To create a calibration curve, pHluorin expressing cells were grown to midlogarithmic phase in YPD at 37°C, and 150 μl were pelleted in 8 microfuge tubes, washed with sterile deionized water, and resuspended in calibration buffer (50 mM MES, 50 mM HEPES, 50 mM KCl, 50 mM NaCl, 200 mM ammonium acetate) ranging from pH 5 to pH 8.5, in 0.5 increments adjusted with HCl or NaOH. 150 μL of unlabeled cells were also resuspended in calibration buffer to allow subtraction of autofluorescence. Cells were assessed for median fluorescence intensity using the 405nm laser, with 525/50 BP filter (AmCyan) and the 488nm laser with 530/30 BP filter (FITC) on a BD Fortessa flow cytometer. AmCyan and FITC fluorescence of cells in calibration buffer at each pH was adjusted by subtracting respective fluorescence of unlabeled cells. The emission ratio of 405_ex_/488 _ex_ for each pH was plotted, and the standard curve was interpolated by 3^rd^ order polynomial regression using Graphpad Prism (v10). The 405_ex_/488 _ex_ ratio was determined for pHluorin-expressing cells after subtracting fluorescence values from unlabeled cells, and the standard curve was used to determine pH of samples. For experimental conditions, the 405_ex_/488 _ex_ ratio for pHluorin-expressing cells grown in YPD at 37°C was determined after subtracting fluorescence values from unlabeled WT or *yck2*Δ cells grown in parallel, and pH was determined from the calibration curve.

### RNA-sequencing

WT and *yck2*Δ strains were grown to midlogarithmic phase in YPD media at 30°C, and control samples were collected. The remaining cells were resuspended in pre-warmed 37°C media for one hour, then pelleted and flash frozen. RNA was extracted by mechanical disruption using glass beads in RLT buffer with 1% β-mercaptoethanol. Samples were then purified using the Qiagen RNeasy kit, following manufacturer instructions including on-column DNase I digestion. rRNA integrity was visualized by gel electrophoresis, then samples were sent for library preparation, poly(A) purification, and RNA sequencing on an Illumina platform (Azenta). 350M paired-end reads were collected (2 x 150bp). Two biological replicates were sequenced for each sample.

#### RNA-seq analysis

FASTQ files were processed to generate a counts matrix using the following software in command-line: Cutadapt/3.2 (adapter trimming and quality control), STAR/2.7.2b (alignment), and RSEM/1.2.20 (read counting) (53–55). The programs FastQC and multiQC were used along this pipeline for quality control and to monitor alignment. Cutadapt was run with quality filter of 10, and minimum read length of 1 to remove empty reads and was specified to trim the Illumina universal adapter. STAR was run in read alignment mode with default parameters using the FungiDB H99 genome build (version 63) (56). RSEM was run with a forward probability of 0.5, again using the FungiDB H99 genome build (v63) (56). All subsequent analysis was performed in R.

For differential expression, read counts were rounded and analyzed in pair-wise fashion using DESeq2 (57). The resulting differential expression tables were then filtered for an absolute value of log2(fold-change) of at least log_2_(1.75), and p-adjusted value < 0.05. Transcripts meeting these thresholds were considered differentially expressed for all analyses.

#### Visualization of RNA-seq data

Visualization of RNA-seq data was performed in R (58).

#### Volcano plots

Volcano plots for pairwise comparisons were generated using the R package EnhancedVolcano (59), with custom gene lists imported and colored as indicated. For volcano plots highlighting transcripts under specific GO terms, ontologies were obtained from FungiDB (v68) (56). The genes corresponding to “oxidoreductases” and “RP transcripts” are stored under the identifiers GO:0016491 and GO:0003735. Dashed lines in volcano plots represent the significance and fold-change cutoffs described above for differential expression.

#### GO analyses

Bubble plots were generated using ggplot2 (60) based on GO analyses performed using the gene ontology tool on FungiDB (v68) (56). Point size represents the log10(p-value), and the x-axis shows counts and is log-scaled. Points were colored based on the ontology from which they originate.

### In vitro macrophage uptake assays

THP-1 monocytes were differentiated to macrophages by incubation in RPMI complete media supplemented with 50 nM phorbol 12-myristate 13-acetate (PMA) for 48 hours, followed by rest for 24 hours. *C. neoformans* Pma1mNeonGreen or Pma1mNeonGreen:*yck2*Δstrains were grown overnight in YPD at 30°C or 37°C, washed in PBS three times, and resuspended in PBS. To opsonize *C. neoformans*, cells were incubated with baby rabbit complement (10%) at 37°C for one hour. Opsonized or unopsonized *C. neoformans* cells were added to macrophages at MOI = 5 in reduced serum media (RPMI + 1% FBS) and incubated for 2 hours at 37°C, 5% CO_2_. Cells were washed 4 times with cold PBS. Cells were lifted with TrypLE for 20 minutes at 37°C. Samples were washed in cold RPMI, incubated with Fc block for 10 minutes, followed by staining with anti-huCD45:PerCP-Cy5.5 (Invitrogen, clone HI30). Cells were fixed in 2% para-formaldehyde for 10 minutes. Cells were assessed for uptake by flow cytometry on a BD Fortessa cell cytometer, and data was analyzed using FlowJo v10. Macrophages that phagocytosed *C. neoformans* were identified by gating for singlets, CD45^+^, and then mNG^+^ cells within the CD45^+^ population.

### Mouse infection and bronchoalveolar lavage

Overnight grown Pma1mNeonGreen or Pma1mNeonGreen:*yck2*Δ cells (YPD, 30°C) were pelleted, washed in PBS, and diluted to 10^8^ cells/ml in PBS. Nine 5-7 week old female BalbC/J mice per group were anesthetized with isoflurane and intranasally infected with 25 μL of inoculum (2.5 x 10^6^ cells). 3 naïve mice were included as controls. After 24 hours, mice were sacrificed by asphyxiation with CO_2_. To obtain bronchoalveolar lavage (BAL) a small incision was made in the trachea and a 20G blunt end needle was inserted and secured tightly with dental floss to prevent leakage. 1.5 mLs of PBS was gently pushed into the needle with a syringe, inflating the lungs, followed by gently suctioning the fluid. Plunging and suctioning was repeated twice more and the final fluid retrieved was collected for immediate processing for flow cytometry. All animal work was performed in compliance with IACUC protocol MIC 37048Y.

### Flow cytometry of BAL cells (Eric)

The BAL was centrifuged for 7 minutes at 400 x g, 4°C. The supernatant was frozen at -80°C. The pellet was treated with 200 uL of ACK Lysis Buffer for 2 minutes at room temperature to lyse red blood cells followed by the addition of 1 mL of cold PBS. Cells were collected by centrifugation (400 x g, 7 min, 4°C), washed with 100 uL PBS, then resuspended in 65 μL of the cell staining solution for 20 minutes on ice. The cell staining solution included Live/Dead fixable Blue Dead Cell Stain (Invitrogen), anti-CD45-APC-ef780 (Invitrogen, clone 30-F11), anti-CD11b-BV605 (BD Horizon, clone M1/70), anti-F4/80-PE-Cy7(Invitrogen, clone BM8), anti-CCR2-PE (RD Systems, clone FAB5538P), and anti-Ly6G-PE-CF594 (BD Horizon, clone 1A8) with Fc Block (BD Horizon, clone 2.4G2) and mouse serum prepared in HBSS. Cells were fixed in 2% para-formaldehyde for 10 minutes. Cells were assessed for uptake by flow cytometry on a BD Fortessa cell cytometer, and data was analyzed using FlowJo v10. The gating scheme for cell populations can be found in Supplementary Figure S7.

### Quantification and statistical analysis

Statistical analyses were performed using GraphPad Prism (version 6.05) software. For all analyses, significance was defined as follows: \**p* < 0.05; \*\**p* < 0.01; \*\*\**p* < 0.001. Specific statistical tests for individual experiments can be found in figure legends.

## Data Availability

RNA sequencing data for *C. neoformans* WT and *yck2*Δ have been deposited in the GEO under code GSE320044. The mass spectrometry proteomics data have been deposited to the ProteomeXchange Consortium via the PRIDE (61) partner repository with the dataset identifier PXD083302.

All other relevant data are available from the corresponding author on request.

## Supporting Information

All supplementary tables can be found in Supporting Information_S1.

**S1 Fig. Signatures of translational reprogramming are intact in the *yck2*Δ mutant.** A. Repression of *RPL2* following a shift from 30C to 37C was determined by northern blot, *n* = 3. B. Activation of Gcn2 following a shift from 30°C to 37°C was assessed by western blot detecting phosphorylation of eIF2α. Images are representative of 3 biological replicates. Polysome profiles for WT (C) and *yck2*Δ cells (D) were recorded for cells grown to midlogarithmic phase at 30°C and following a shift to 37°C for one hour. Images are representative of 3 biological replicates.

**S2. Fig. Levels of chitin and exposed chitooligomers are elevated in the absence of Yck2.** Cells were grown to midlogarithmic phase at 30°C or 37°C and stained for chitin with calcofluor white (CFW) and exposed chitooligomers with wheat-germ agglutinin (WGA). Cells were imaged by fluorescence microscopy. Scale bar = 10 μm, images are representative of 3 biological replicates.

**S3 Fig. Levels of cell wall chitosan are elevated in the absence of Yck2.** Cells were grown to midlogarithmic phase at 30°C or 37°C and stained for chitosan with eosin Y. Cells were imaged by fluorescence microscopy. Scale bar = 10 μm, images are representative of 3 biological replicates.

**S4 Fig. Absence of Pdr15 or Tpo3 do not affect antifungal susceptibility.** The *yck2*Δ, *YCK2*:*yck2*Δ, and *pdr15*Δ mutants and their parental background, H99, and the *tpo3*Δ mutant and its parental KN99 background were spotted onto YPD plates containing indicated antifungals. Plates were incubated at 30°C for 3 days and photographed. Images are representative of 2 biological replicates.

**S5 Fig. Ribosomal protein transcript repression occurs in the absence of Yck2 in response to host temperature stress.** Volcano plots display DEGs in response to host temperature stress in the WT (left) and yck2 mutant (right). Ribosomal protein (RP) transcripts are highlighted in red.

**S6 Fig. *C. neoformans* Pma1 lacks the conserved Yck1/2 phosphorylation site and has an extended C-terminal cytoplasmic tail.** A. Sequence alignment of the alpha helix in the N domain region of *S. cerevisiae* Pma1 (top) and *C. neoformans* Pma1 (bottom) that harbors the Yck1/2 phosphorylation site in *S. cerevisiae* (indicated in blue). B. The *C. neoformans* Pma1 (cyan) was modeled on top of *S. cerevisiae* Pma1 (yellow). Serines and threonines in the C-terminal tails are highlighted in magenta and blue, respectively.

